# Implantable Electrical Connector for Reconnecting Injured Neurons, Approach and Design

**DOI:** 10.1101/2020.05.29.123232

**Authors:** Soheil Hashemi, Ali Abdolali

## Abstract

In contrast to normal injuries, damages of the nervous system are very difficult to repair. In this paper, a connector is designed to reconnect injured nerve to healthy cells. This cord was designed and tested by the full wave and three dimensional electromagnetic modeling. The connector is comprised of three parts, including the conductor, dielectric coating, and second coating with high conductivity. Electromagnetic characteristics need to be unique to work fine, each part is obtained with regard to the distance between two neurons. The combination of materials with conductivity lower than 100 S/m and relative permittivity lower than 8 creates the connection of two neurons up to 35 cm. The connector is realizable by biocompatible polymers.

## I. INTRODUCTION

In contrast to normal injuries in body, damages of the human nervous system are very difficult to repair and this system has limited repairability[1]. In small injuries, a damaged cell would grow toward the ruptured part and generate the connection[2-4]; however, this would be impossible if the cutting exceeded a certain limit and consequently, surgery would be necessary[5]. If the cut is sharp and the cell is not destroyed, the two ends get closer to create the connection[6,7]; otherwise, a bridge is used to lead the cell in right direction and establish the connection gradually[8-11]. If this is not the case, a cell should be selected from an unimportant part of the body and replaced with the damaged cell^1^. However, all of these methods are not practical in certain circumstances.

This capability becomes more limited in the central nervous system compared to the peripheral nervous system and is stopped by glial cells when regeneration of the cells is initiated[1,12]; thus, it is always necessary to use novel methods for connecting or regenerating damaged nerve cells. Such necessity is intensified when the cell is destroyed, or impossible to be repaired. Hence, the present paper proposes a method for connecting a healthy and damaged cell with no possibility of growing to another cell. On this basis, a cord with physical structure and the designed material is proposed which goes from the first cell to the second cell and transmits the action potential. This structure is designed by simulation, and resulted diagrams of the design are presented in following sections. Specifications of the material and structure differ depending on the distance that should be connected, which will be discussed later.

## II. Method

Two possibilities commonly emerge in complete injuries. first, one neuron is healthy and the other is irreparably damaged, and thus it is necessary to develop connection and signal transmission between these two cells; second, a neuron is destroyed; thus, two healthy cells are existed but there is a long distance between them, for which a connection is required to be established.

In order to solve this problem, it is proposed to consider a cord as shown in Figure (1). This cord is structured as three parts: cord’s conductor and two coating layers. In case each end of the cord’s conductor part is connected to the first and second cell, signal propagation from first cell to the second cell will be possible through the proposed connector. The conductor part should be in contact with intracellular fluid of both the cells and must never have contact with extracellular fluid; therefore, it is necessary to use a coating with very low electrical conductivity to cover the cord’s conductor. In order to prevent the cord from changing the cell function while propagating the signal, the material and diameter of the cord’s conductor and coating layer must be specified exactly. Material and diameter of the cord’s components depend on the distance between the two cells and approximate diameter of the cells. Rate of the potential change in the cord’s conductor should be adjusted such that the propagated signal can reach the next cell and, meanwhile, does not cause unwanted stimulation, and thus the cells can achieve stability in resting state. It is seen in Figure (1) that a layer with electrical conductivity of 10^5^ has been placed on the connector to reduce the effect of peripheral noises on the cord.

**Fig. 1.**
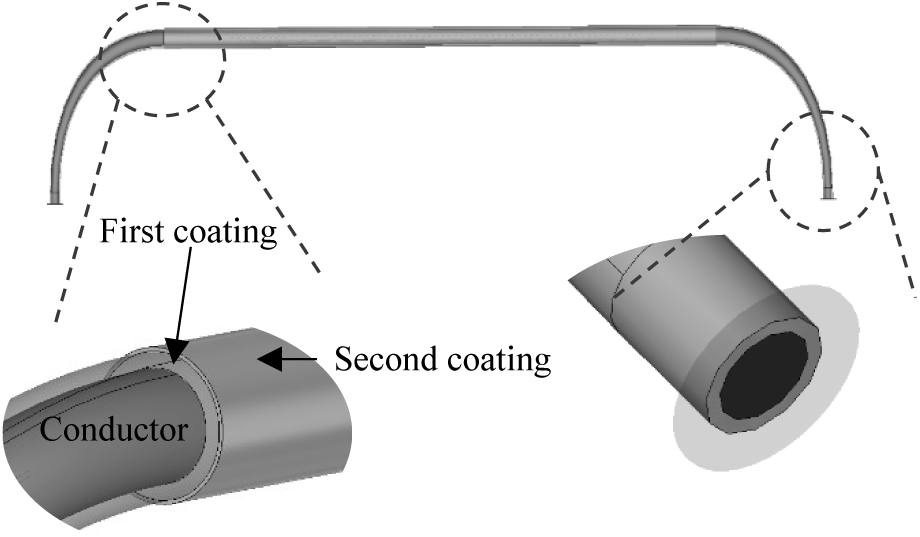
Designed connector comprised of conductor parts with high electrical conductivity and a dielectric coating layer with very low electrical conductivity. Second coating with high electrical conductivity on the first coating with low electrical conductivity; this layer secures the conductor part and eventually, the connected cells against the external electromagnetic noises.

To find the appropriate characteristics of each part which satisfy mentioned conditions, this connection was simulated 3-dimensionally between the two neurons using electromagnetic modeling, as shown in Figure (2). We propose equations (1) to (2) for 3D modeling of neurons. The ionic currents and fluxes varying with time were inserted in Maxwell’s equations as the source current densities[13-16]:

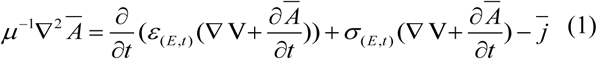

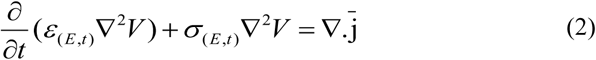

where *A, B, V, E*, and *j* indicate the magnetic vector potential, magnetic field flux, electric potential, electric field, and electric current density, respectively. *ε* is the permittivity and *σ* is the conductivity.

**Fig. 2.**
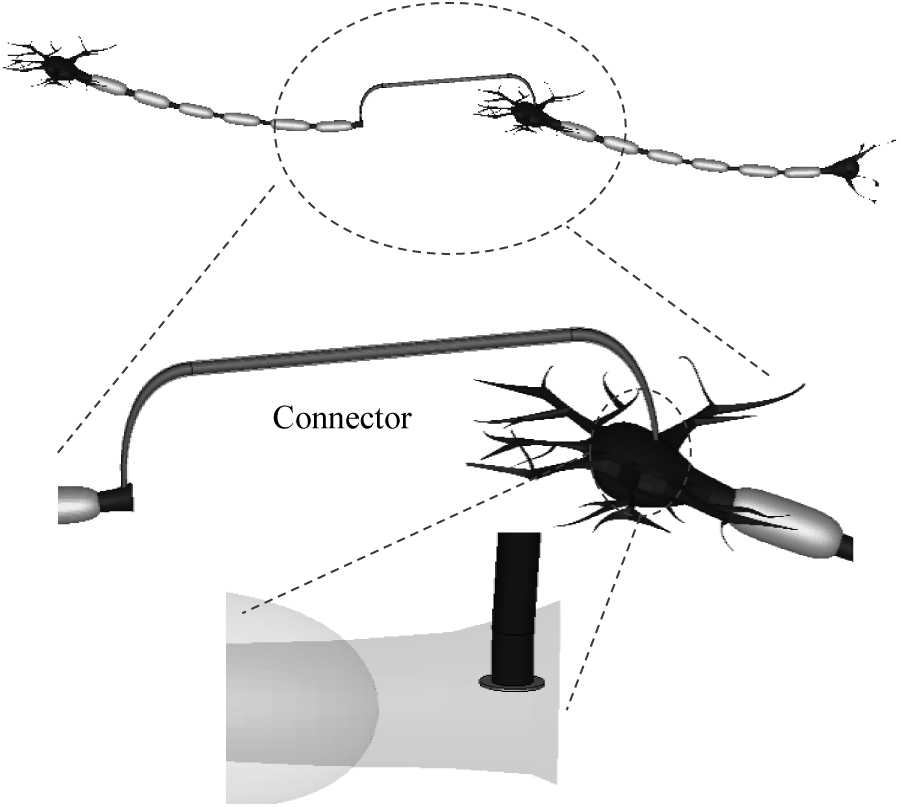
The way of connecting a nerve cell to the ruptured axon terminal to a healthy nerve cell by the connector. Conductor part of the connector enters the intracellular fluids of both the cells and is isolated from the extracellular fluid by the outer coating.

Using the partial differential method, we carry out discretion of equation 1 and 2. Forward methods are used commonly in time domain though stability conditions of forward methods force us to select small time step due to dimensions of neurons which takes huge amount of time. To avoid this issue and solve these equations versus time, we propose using backward method and carrying out their discretion to obtain unknowns according to known variables in last time step. The discrete equations of 4 and 5 by considering inhomogeneity of the surrounded media are shown in 3 to 6.

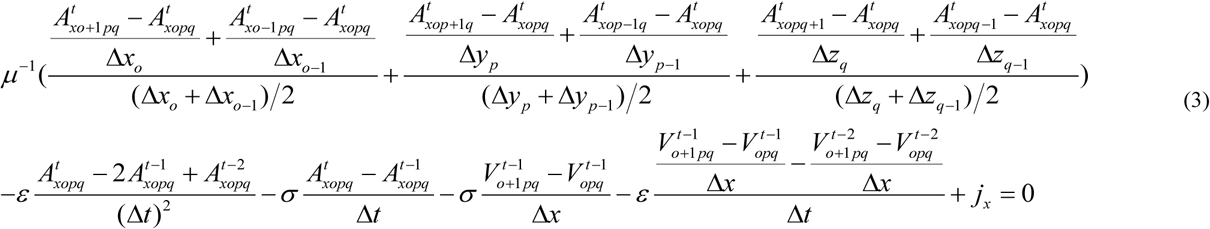

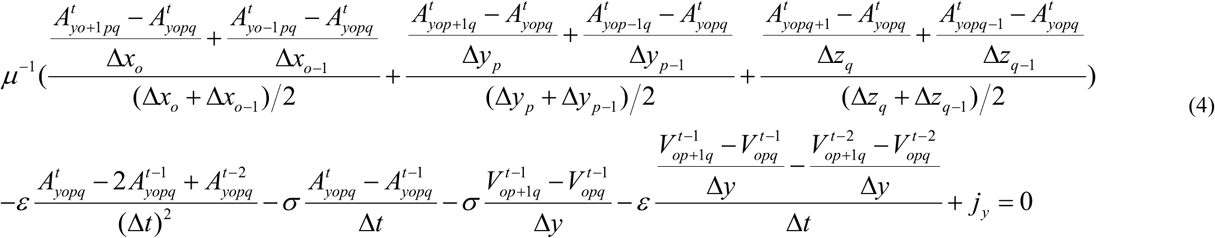

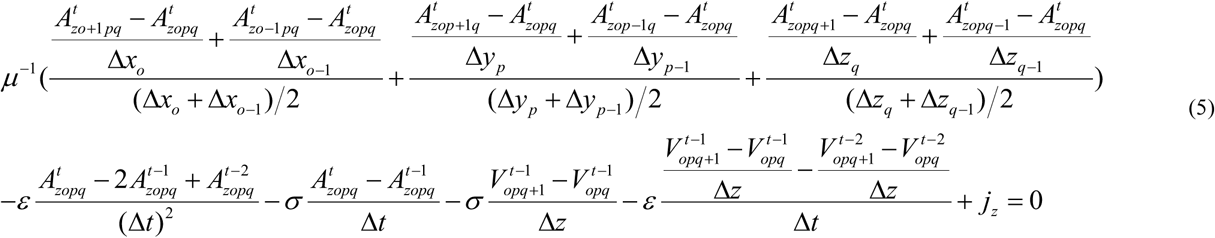

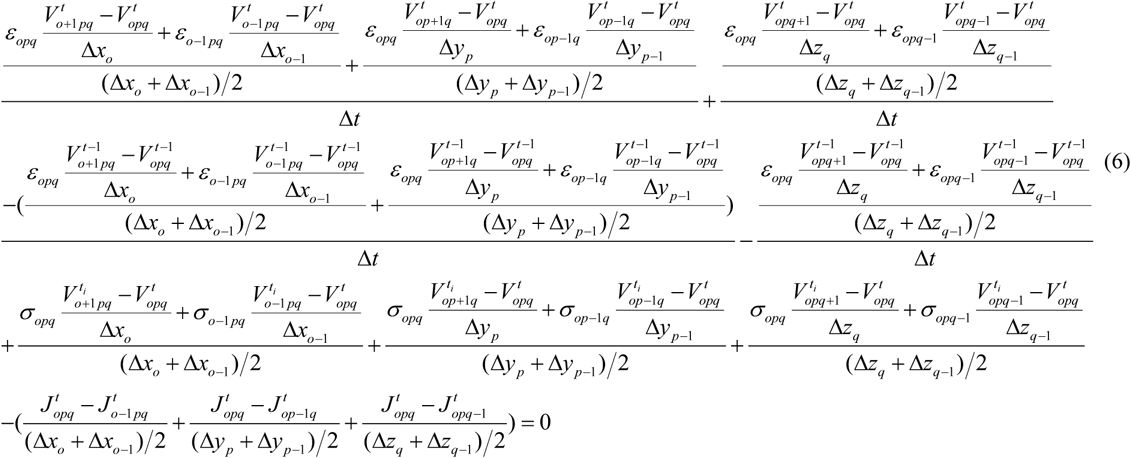

Where Δ*x*, Δ*y*, Δ*z* are discrete distances in each axis. o, p, and q indicate the number of discrete points considered at directions x, y, and z. Superscript t shows the time step. Subscripts x, y, and z present vector potential *A* on those directions. After the discretion a system of linear equation appears in each time step as 7, which does not converge by common recursive method.

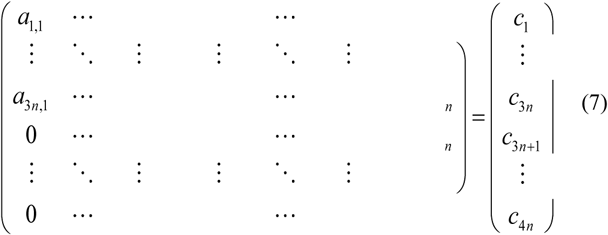

Where *n* equals to multiplication of the number of discrete points considered at directions *x, y*, and *z*. We have used methods for sparse matrices to solve the equation and find unknowns in each time step.

In low frequencies analysis, Perfect Electric Conductor (PEC) is commonly used as the boundary conditions. Hence, we have also used the PEC boundary condition (*V* is constant, *At*=0, and 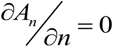) on the boundaries. Electrical conductivity of the cells’ membrane was considered as varying with time, as in Hodgkin and Huxley equations in which it varies with time and voltage. The used equations, which control the currents and electrical conductivity in mammals, can be observed in McIntyre’s works[17,18]. By investigating Maxwell equations and cable model as 8, we propose to obtain the neuron characteristics required in the equations 3 to 6 from the properties mentioned in cable method based on the following equations.

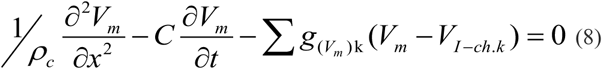

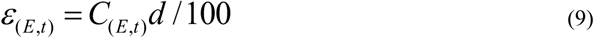

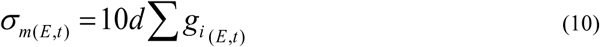

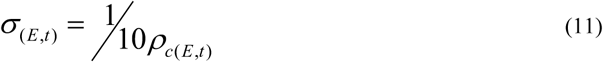

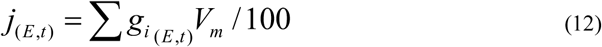

Where *d* is the membrane diameter, *g* is membrane ions conductivity, C is the capacitance of the membrane in the cable model, *ρ*_*c*_ is circuit model series branch resistance or the longitudinal resistance of the cell, and *Vm* is membrane voltage in the cable model. Using this method, a three-dimensional (3D) cell can be assumed and the action potential (AP) propagation in the simulated neuron is achieved similar to a real neuron. The proposed method is validated by literature [19-20]. The required structure, dimensions and characteristics of the connector are obtained using the simulations by the proposed method.

## III. Results and discussion

The simulation results of the propagation of an action potential from an injured cell to another cell through the designed connector is shown in Figure (3). As can be seen, the action potential in the first stimulated cell has been propagated to its end and then to the second cell using the cord; subsequently, the action potential has been stimulated and propagated in the second cell. Conductivity, permittivity and diameter of each of three parts of the connector should be chosen simultaneously. We have shown the characteristics required for connecting two cells in accordance to their distance in Figure (4).

**Fig. 3.**
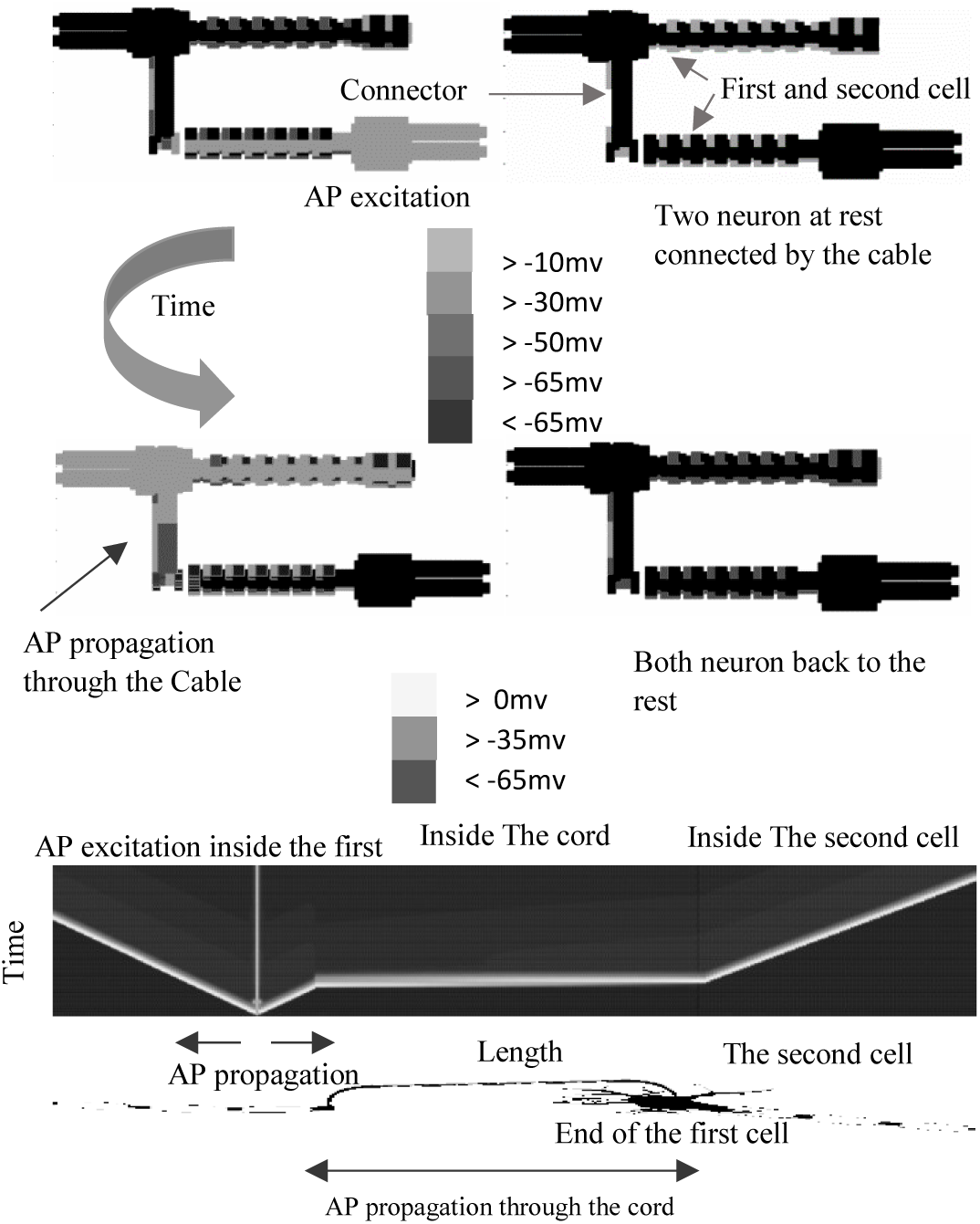
Stimulation and propagation of action potential in a cell and its transmission to the next cell by the designed connector. Length and radius of the connector are 12cm and 1μm, respectively, conductivity of the cord’s conductor is 30S/m, diameter and relative electrical permittivity of coating are 1 μm and 5, respectively, and Soma’s radius of the nerve cell is 3μm.

**Fig. 4.**
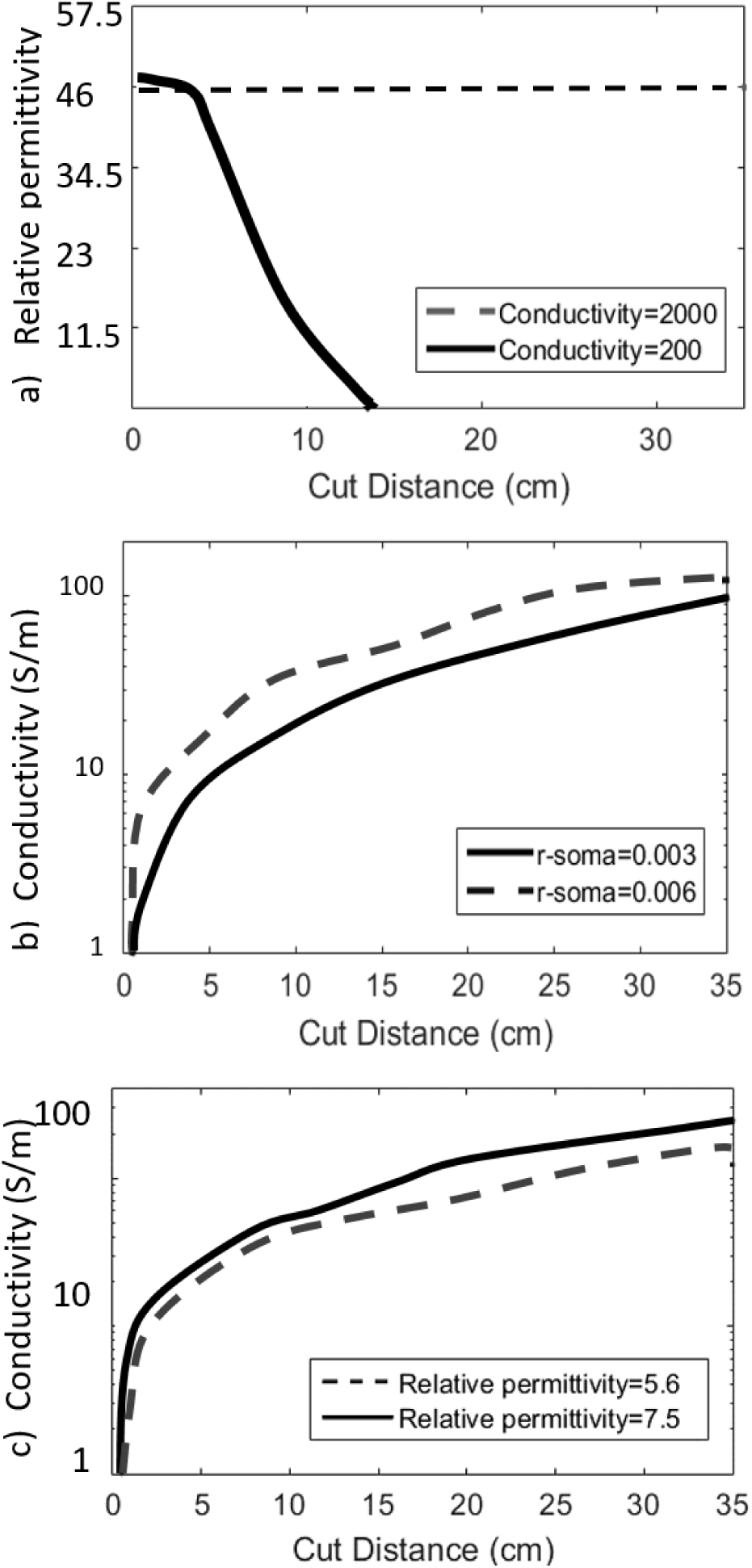
Electromagnetic characteristics of the connector in accordance with the required distance between two neurons. a) The maximum electrical relative permittivity required for a coating layer with diameter of 1μm on the cord’s conductor part with radius of 1μm and electrical conductivity of 200 and 2000S/m; b) The minimum electrical conductivity required for the cord’s conductor part in two cases of connection between two cells with approximate Soma’s radius of 3 or 6μm, in case of coating with a layer with thickness of 1μm and electrical relative permittivity of 11.3; c) The minimum electrical conductivity required for the cord’s conductor part in two different coating cases with electrical relative permittivity of 5.6 or 7.5, in case of coating with a layer with thickness of 1μm and approximate cellular Soma’s radius of 3μm.

Different scenarios is shown, in Figure (4-a) the conductance of the conductor is set at two cases of 200 and 2000 S/m and required relative permittivity of the first coating is shown. This diagram is resulted for coating with 1 μm thickness, by decreasing the thickness lower relative permittivity is required to decrease existence capacitance between outside liquid and conductor of the connector. The limit for decreasing the thickness is achieved when the required of relative permittivity of the first coating decreased to one, for example when conductivity of internal conductor of the cord is 2000 S/m, the minimum thickness of first coating is 20nm. Figure (4-b) illustrate required conductivity of internal conductor of the cord for two different nerve fiber with 3 and 6 μm diameter. By increasing the diameter of the nerve fiber thicker conductor and more conductivity are required for the propagating AP between the fibers. In Figure (4-c), required conductance is illustrated on two cases of first coating. It shows that by increasing relative permittivity of the first coating, more conductance is needed on the internal conductor and by increasing distance between two nerves, more conductance is required.

Based on the simulations, increased diameter and decreased electrical conductivity of the coating caused better signal propagation and higher stability of the cell function; on the other hand, increased diameter and electrical conductivity of the cord’s conductor also led to better signal propagation.

Characteristics of the coating and conductor part should be realized simultaneously in order for propagation to be accomplished; this can be seen in Figure (4-a). For the state of the 200S/m electrical conductivity of the cord’s conductor, propagation will not be accomplished howsoever the coating permittivity is reduced.

Dimensions and characteristics mentioned in diagrams of Figure (4) are constructible using the biocompatible polymers. To examine these polymers, the specifications noted in the background history of these polymers can be investigated [21,23]. Polythiophene or Polypyrrole polymers and Emaraldine base PANI polymers can be used for constructing the cord’s conductor and coating, respectively. Using these diagrams, the required characteristics can be chosen with regard to the distance between the two disconnected cells and diameter of the cells, and also the corresponding cord can be used for connecting the two cells. In the literature many methods are proposed for healing the injured neve fiber, and repairing the nerve connections such as use of bridges, surgeries or replacing the nerve [1,11]. In this paper, a new method by using a designed cord for repairing connection between injured nerves is proposed and discussed even when other methods are inefficient.

## IV. Conclusions

Using the proposed cord, injured neurons can be replaced and reconnected. Distances of up to 35cm have been investigated in the article by simulations where they are proven to be connectable. This device can play an effective role in the spinal cord injuries and other anesthetic diseases. In previous methods, in which a bridge was used for conduction and regeneration of the cell, a limited length was performable only if the cell had not been completely destroyed. In case the cell had been destroyed, another cell should be selected from another part of the body and replaced, which per se resulted in anesthesia in the place from which the cell had been selected. However, in the present method, any disconnection at any distance can be reconnected by this cord; so that, two nerve cells should be selected and then be connected using this connector.

